# Microtubule lattice conformation and integrity regulate α-tubulin acetylation

**DOI:** 10.1101/2025.09.09.675099

**Authors:** Cornelia Egoldt, Marie-Claire Velluz, Joshua Tran, Charlotte Aumeier

## Abstract

Microtubule acetylation of lysine 40 of α-tubulin is a hallmark of stable microtubules. This luminal modification is catalyzed by α-tubulin acetyltransferase 1 (αTAT1) and reversed by histone deacetylase 6 (HDAC6). However, acetylation regulation within the microtubule lumen and the influence of lattice architecture on enzymatic activity remain poorly understood. Here, we reconstitute microtubule acetylation *in vitro* using purified αTAT1 and HDAC6 on microtubules assembled with defined lattice conformations. We show that αTAT1 overwrites HDAC6 enzymatic activity, but its acetylation efficiency decreases upon microtubule damage. Importantly, αTAT1 efficiently acetylates microtubules in expanded lattices and twisted tubulin states, while compacted lattices impede its activity. Our findings reveal that both microtubule integrity and lattice conformation are critical regulators for αTAT1 enzymatic activity, suggesting that dynamic lattice transitions modulate the acetylation pattern of microtubules in cells.

## INTRODUCTION

Microtubules are dynamic cytoskeletal filaments that play key roles for cell shape, motility, and division^1^. Heterodimers of α- and β-tubulin binding guanosine triphosphate (GTP) assemble into polarized microtubule biopolymers, a hollow tube composed mainly of 13 protofilaments (pfs) in eukaryotic cells^2,3^. Within the microtubule lattice, tubulin can adopt different states: a guanosine diphosphate (GDP)-bound, compacted state lattice and an expanded, GTP-bound state at the microtubule tip. The compacted GDP-microtubule lattice is under more strain, with an average dimer rise of 81.5 Å within a protofilament, compared to an expanded dimer conformation of 83.2 Å in the GTP-cap^4–8^.

The level of lattice expansion can be read and/or modulated by different microtubule associated proteins (MAPs): EB3 or DCX bind to the microtubule surface at the interface of 4 tubulin dimers^9,10^, TPX2 binds across longitudinal and lateral protofilaments^11^ and MAP7 and tau bind longitudinally within a protofilament^12,13^. Thereby they reshape the biochemical landscape of microtubules, locally fine-tuning microtubule function and stability.

*In vitro,* by replacing the nucleotide GTP with its slowly hydrolysable analogue guanylyl-(α,β)- methylene-diphosphate (GMPCPP), stable microtubules nucleate with 14 protofilaments in an expanded lattice state (83.2 Å)^4,6,8,14^. In addition, specific lattice conformational states can be generated *in vitro* by using different microtubule stabilizing agents (MSAs). Two distinct ligand-binding sites on β-tubulin exist for MSAs, the taxane site and the laulimalide/peloruside site, both preventing microtubule depolymerization^15^.

Paclitaxel (taxol) has been shown to expand the microtubule lattice inducing a GTP-like state (82.3 Å), with a preference for 13-pf microtubules^4,5,16,17^. The taxane site is in the microtubule lumen and taxol binding has been reported to stabilize β-tubulin in a conformation that favors lateral contacts^18–20^. Furthermore, simulations showed that taxol exerts a “pseudo” kinetic stabilization through decreasing the strain energy of the GDP-tubulin lattice^21^. The laulimalide/peloruside site is located on the outside of the microtubule near the lateral interface of protofilaments^15^. Structural analysis showed that peloruside A acts on heterologous lateral tubulin contacts at the microtubule lattice seam, a discontinuity in the microtubule lattice^5,22^. Through binding to β-tubulin it supports a homologous, non-seam interaction. Opposite to taxol, peloruside A does not induce microtubule lattice expansion but rather compacts the lattice (81.0 Å) and even overrides the effects of Taxol binding^5^.

In cells, microtubules are dynamic and exposed to different stresses like mechanical forces, the activity of motor proteins or severing enzymes, all potentially removing tubulin dimers from the shaft^23–26^. Dimer dissociation can occur spontaneously along the microtubule shaft, as well as at a switch in protofilament number or at a transition in the helix pitch^27,28^. Changes in seam number and seam location can also leave holes in the lattice during microtubule assembly^28,29^. This damage can be repaired to rescue microtubule growth by the incorporation of free GTP-tubulin into the lattice^30–32^.

Previous results from our group and others showed that the processive motor protein kinesin-1 induces shaft damage^24,33,34^. Kinesin-1 motors generate force while running along a protofilament which results in dimer dissociation from the shaft^35–38^. Furthermore, kinesin-1 has been shown to preferentially bind to GTP-tubulin within the lattice, to buckle microtubules and to expand the microtubule lattice^39–42^. A 5-amino acid deletion in the neck linker region results in a kinesin-1 mutant, that damages microtubules more severely while running along them^34^.

In the cellular network, microtubules are highly modified by MAPs which add and remove post-translational modifications (PTMs) to tubulin subunits in a tightly regulated manner. Modifications like de-/acetylation, de-/tyrosination or glutamylation determine the “tubulin code” of specific microtubule subsets^44,45^.

Tubulin acetylation is a modification that occurs on the lysine 40 (K40) residue of α-tubulin which resides in the microtubule lumen upon microtubule polymerization^46,47^. The level of K40 acetylation in the mammalian brain is about 30%^48^. The enzyme α-tubulin acetyl transferase 1 (αTAT1) acetylates tubulin nearly exclusively once it is polymerised into microtubules and must therefore access the lumen for K40 modification^49,50^. Hence, αTAT1 enters through open ends and through damage sites along the microtubule shaft to diffuse and stochastically acetylate α-tubulin^49,51,52^. In cells, this acetylation behavior leads to distinct acetylation segments along a single microtubule^43^. HDAC6 is three-times larger in size (140 kDa) compared to αTAT1 (45 kDa)^50,53^ and deacetylates tubulin with a 1500-fold higher rate than polymerised microtubules^54,55^.

*In vitro* experiments demonstrated that αTAT1 acetylates and HDAC6 deacetylates microtubules stochastically along their entire length^49,54^. In cells, microtubule acetylation forms a gradient, with high acetylation around the nucleus decreasing towards the cell periphery^43^. This gradient results from a tightly controlled interplay between αTAT1 and HDAC6. We recently showed that running kinesin-1- induced microtubule damage, changes the cellular acetylation gradient, reduces cell acetylation levels and results in deacetylation specifically around damage sites^43^. But whether both enzymes counteract each other within the microtubule lumen to modify the K40 residue on α-tubulin and how microtubule lattice conformation impacts enzymatic activity remain unknown.

In this study we test microtubule acetylation in a reconstituted model system. We use purified αTAT1 and HDAC6 proteins on *in vitro* assembled microtubules. We show that αTAT1 outcompetes HDAC6 enzymatic activity but that αTAT1 acetylation efficiency decreases when microtubules are damaged. We demonstrate that not only loss of microtubule integrity but also the state of the underlying microtubule lattice impacts acetylation efficiency, as αTAT1 prefers to acetylate microtubules in an expanded state.

## RESULTS

### αTAT1 overwrites HDAC6 activity on intact microtubules in vitro

Confirming previous findings^49,54^, our *in vitro* reconstitution assay shows that HDAC6 preferentially deacetylates non-polymerized brain tubulin and αTAT1 preferentially acetylates preformed microtubules (GMPCPP-stabilized). Notably, microtubule length does not affect αTAT1 acetylation efficiency^56^. Within 60 min, HDAC6 deacetylates brain tubulin nearly completely, while reducing polymerized microtubule acetylation levels by only 20% compared to control conditions (Fig. 1a,b). αTAT1 on the other hand barely acts on free tubulin but increases microtubule acetylation levels by 5.5-fold within 60 min (Fig. 1a,b).

**Fig. 1.**
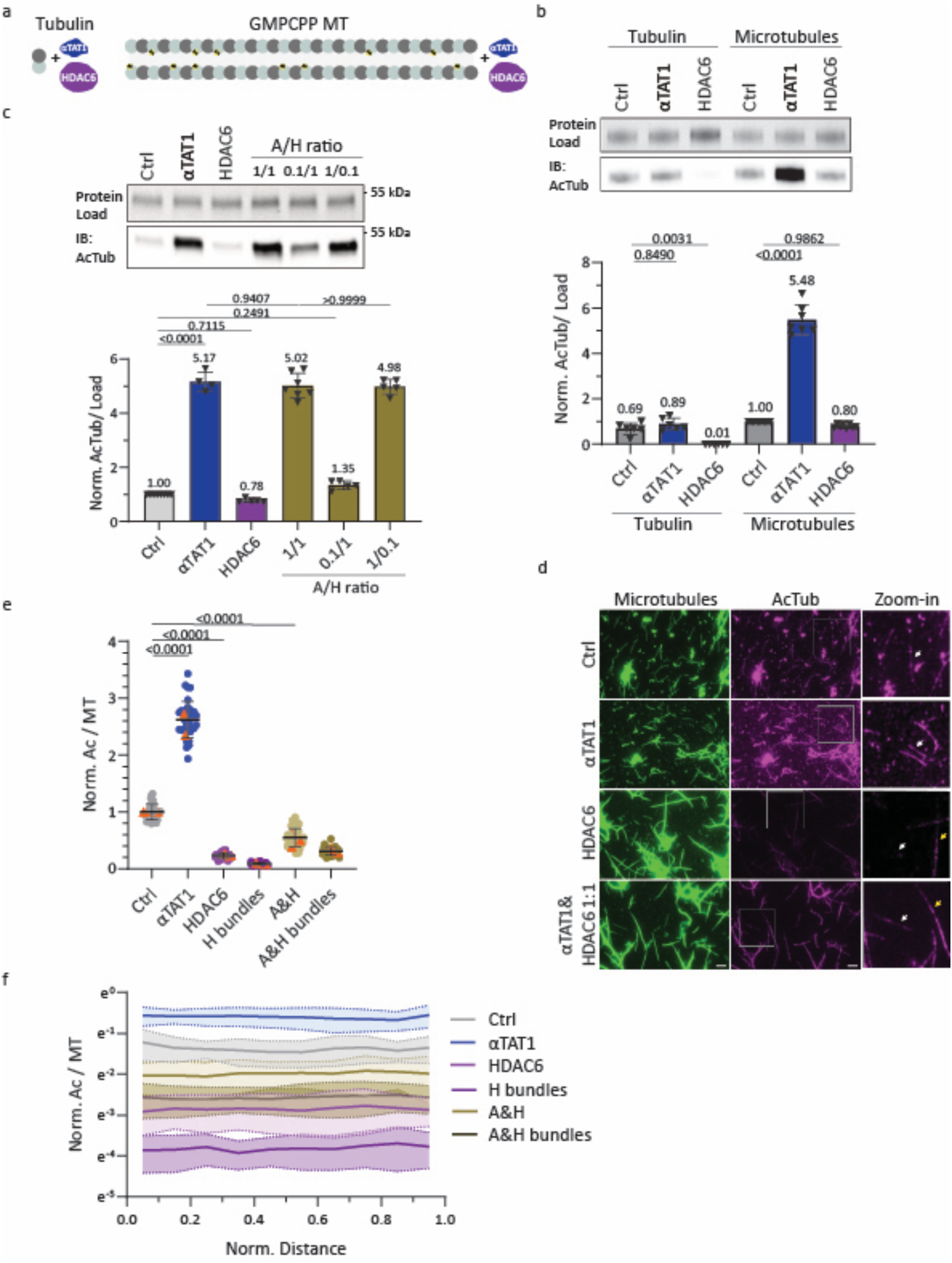
αTAT1 overwrites HDAC6 activity on intact microtubules *in vitro*. **a** Cartoon showing the *in vitro* reconstitution setup for unpolymerized tubulin or GMPCPP-microtubules with αTAT1 and HDAC6. **b** and **c** Representative Western blot analysis for (de)acetylation reactions with (**b**) αTAT1 and HDAC6 in presence of tubulin or GMPCPP-microtubules and (**c**) for GMPCPP-microtubules with αTAT1, HDAC6, or different A/H (αTAT1/HDAC6) ratios. Quantifications of AcTub levels relative to the protein load, from 5-7 (**b**) or 4-7 (**c**) independent experiments. Statistics: one-way ANOVA. Mean with SD. **d** Representative immunofluorescence images of fluorescent microtubules (green) that were antibody stained for acetylated tubulin (magenta); Scale bars: 10 µm. The boxed areas show zoom-ins for the respective acetylation levels, with white arrows pointing at single microtubules and yellow arrows at bundled microtubules. Scale bar: 5 μm. **e** and **f** show normalized acetylation levels over microtubule fluorescence for Ctrl (n=30), αTAT1 (n=33), HDAC6 (n=36), HDAC (H) bundles (n=30), αTAT1 and HDAC6 (A&H 1:1; n=45) or αTAT1 and HDAC6 (A&H) bundles (n=30), from 3-4 independent experiments. In **e** the average acetylation signal (orang dot: experimental mean) per microtubule is shown and in **f** the normalized acetylation signal is plotted over the normalized distance of each microtubule. Statistics: one-way ANOVA. Mean with SD.

When both enzymes are present in the reaction at a 1:1:1 ratio with polymerized tubulin, αTAT1 shades the deacetylation activity of HDAC6 and acetylation increases 5-fold despite the presence of HDAC6 (Fig. 1c). Given that a cell expresses HDAC6 with a roughly 6x fold higher protein copy number compared to αTAT1^57^, we reduced the amount of αTAT1 in the reaction. Lowering the amount of αTAT1 by 10-fold while keeping the HDAC6 and tubulin concentration constant, still pushes the system towards a 1.35-fold acetylation increase compared to control conditions. Lowering HDAC6 by 10-fold while keeping tubulin and αTAT1 stable shows no impact on acetylation efficiency (Fig. 1c). Hence, HDAC6 deacetylation activity is low in microtubules so that the presence of its counteractor αTAT1 overwrites its activity.

To visualize the acetylation level along GMPCPP-stabilized microtubules, we used antibody staining. Immunofluorescence reveals a 2.6-fold increase in acetylation in the presence of αTAT1 (Fig. 1d,e) which is lower but consistent with our Western blot data. In contrast, HDAC6 treatment leads to a deacetylation of 80%, substantially more than observed by Western blot. When both enzymes, αTAT1 and HDAC6, are present at a 1:1:1 ratio with tubulin, the antibody showed a 40% decrease in acetylation whereas Western blotting shows an increase (Fig. 1d,e). These discrepancies suggest that the antibody is less efficient to recognize acetylation levels by immunofluorescence *in vitro* than by Western blot.

In addition, HDAC6 not only acts as a deacetylase but also promotes microtubule bundling (Fig. S1a). Bundled microtubules in HDAC6 or αTAT1/HDAC6 conditions show even lower acetylation signals compared to single microtubules (Fig. 1d,e). Furthermore, the acetylation signal is inhomogeneous when microtubules were treated with HDAC6 resulting in patches along the microtubule shaft (Fig. 1d), consistent with previous findings by Soppina et al^46^. Quantitatively, this patchiness was reflected in a higher standard deviation of fluorescence intensity compared to control or αTAT1-only conditions (Fig. 1f). The acetylation reduction and its patchiness is not due to general steric hindrance for antibodies due to HDAC6 binding to microtubules, as α-tubulin staining remained unaffected (Fig. S1b). However, it is possible that HDAC6 interferes with antibody access to the luminal epitope. Given prior *in vitro* findings showing that taxol disrupts antibody recognition^46^ (Fig. S1c) and considering that additional lattice inhomogeneities (*e.g.* lattice expansion) may further impair this antibody to bind, we decided to base our quantitative analysis primarily on Western blotting.

### Characterization of microtubule damage

In cells, microtubule can get damaged through different mechanisms, like mechanical strain, motor proteins or microtubule severing enzymes^24,25^. To reconstitute microtubule damage *in vitro*, we sought to induce damage artificially using two different methods. First “Snap freezing”, where we performed 5 cycles of snap freezing followed by quick thawing of GMPCPP-stabilized microtubules to induce damage to microtubules (see Methods for more details). Second, to induce more physiological relevant damage, we purified the highly damaging kinesin-1 mutant, kinesin-1Δ6^34^ and incubated it with GMPCPP-stabilized microtubules, as wildtype kinesin-1 damages stabilized microtubules insufficiently (see more details in Methods). To characterize microtubule damages induced by the two methods, we used electron microscopy (EM). We categorized observed damages into four classes of increasing severity: dimer dissociation, kink, sheet opening and broken microtubules (Fig. 2a). Dimer dissociation and kinks typically involve the loss of a few dimers from the lattice, though kinks likely reflect a more extensive local disruption. Sheet openings, by contrast, can extend from 100 nm up to 2 µm in length.

**Fig. 2.**
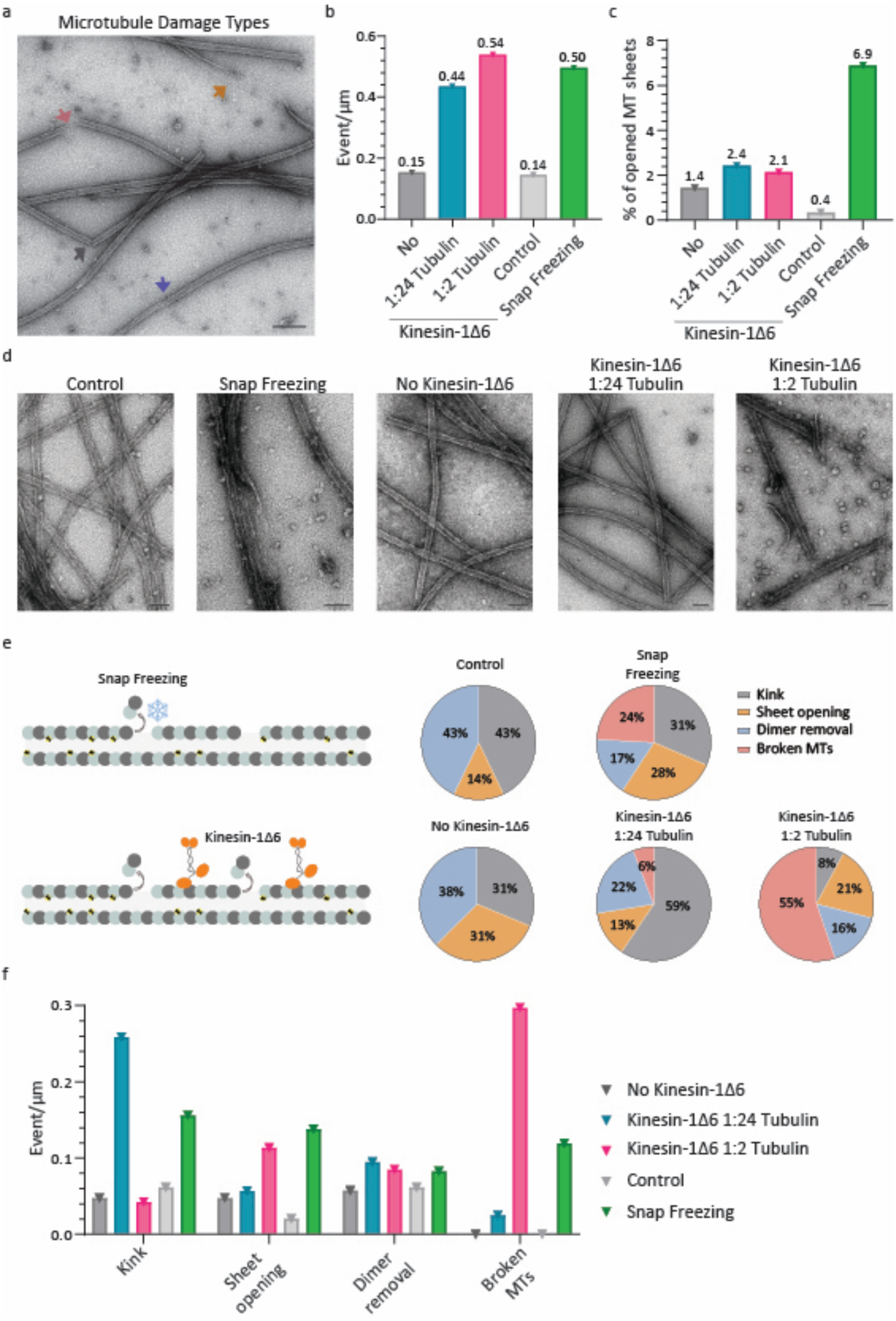
Diversity and abundance of microtubule damage. **a** Representative Electron microscope (EM) image of different damage types of GMPCPP-stabilized microtubules incubated with kinesin-1Δ6 in a 1:24 tubulin ratio. The orange arrow points at a “sheet opening”, the light coral arrow at a “broken microtubule (MT)”, the gray arrow at a “kink” in the microtubule and the blue arrow at a “dimer removal” along the shaft. Scale bar: 200 nm.**b** Damage frequency plotted as event per micrometer along microtubules for both damaging conditions. **c** Percentage of microtubule in sheet opening state observed along microtubules by EM for both damaging conditions. **d** Representative EM images for different damaging conditions. **e** Cartoons displaying microtubule damage generation *in vitro* per either snap freezing or running kinesin-1Δ6-induced damage (left; more details see Methods). Distribution (%) of different damage types per condition (right). **f** Damage frequency as event per micrometer of different damage types for all damaging conditions.

Under control conditions, microtubules were damaged with a frequency of 0.14 events per micrometer (Fig. 2b). Among these, 43% were kinks, 43% sheet openings, and 14% were dimer dissociation along the lattice (Fig. 2e). Note that microtubule damage has previously been reported using various techniques^30,58–60^ but could be amplified by the staining procedure on the EM grid (Fig. 2a, see Methods).

Snap freezing significantly increased the incidence and severity of lattice damage. Microtubules were affected, with a frequency of 0.5 events per micrometer (Fig. 2b). Notably, sheet openings became more extensive, reaching up to 2 µm in length, and their occurrence rose by 17-fold, compared to controls (Fig. 2c). Upon snap freezing, all four damage types were observed, including broken microtubules and long sheet openings (Fig. 2d-f, S2a).

Kinesin-1Δ6 treatment at a 1:2 kinesin:tubulin ratio also induced substantial microtubule damage with a frequency of 0.54 events per micrometer (Fig. 2b); however, unlike snap freezing, the predominant damage was broken microtubules, accounting for 55% of all observed events - indicating a qualitatively distinct damage profile (Fig. 2d-f). At a lower kinesin-1Δ6 concentration (1:24 ratio), the motor primarily induced kink-type damage, which made up 59% of the total damage (Fig. 2d-f). This suggests that at higher concentrations, kinesin-1Δ6 breaks microtubules by first generating lattice kinks that eventually lead to rupture (Fig. 2a, d). Sheet opening, however, only increases 1.5-fold upon motor running on microtubules, which were also shorter, ranging between 100-900 nm, compared to snap freezing induced openings (Fig. 2c). Together, these findings establish two distinct approaches that produce robust yet qualitatively different forms of microtubule lattice damage: snap freezing, which favors extensive sheet openings, and kinesin-1Δ6, which predominantly generates kinks and breakage.

### Microtubule damage reduces αTAT1 acetylation activity and increases HDAC6 deacetylation

We next tested how damage in general - and how these different damage types in particular - affect the enzymatic efficiency of αTAT1 or HDAC6. HDAC6 efficiency to deacetylate kinesin-1Δ6 damaged microtubules is comparable to intact microtubules (Fig. 3b). In contrast, HDAC6 showed markedly higher activity on snap-frozen microtubules, resulting in an 80% reduction of brain tubulin acetylation levels. This is much closer to the enzyme’s efficiency on free tubulin and is significantly greater than the 20% reduction observed in undamaged microtubules (Fig. 3a). These results suggest that the large sheet openings caused by snap freezing enhance HDAC6 access to the luminal K40 site (Fig. 3c).

**Fig. 3.**
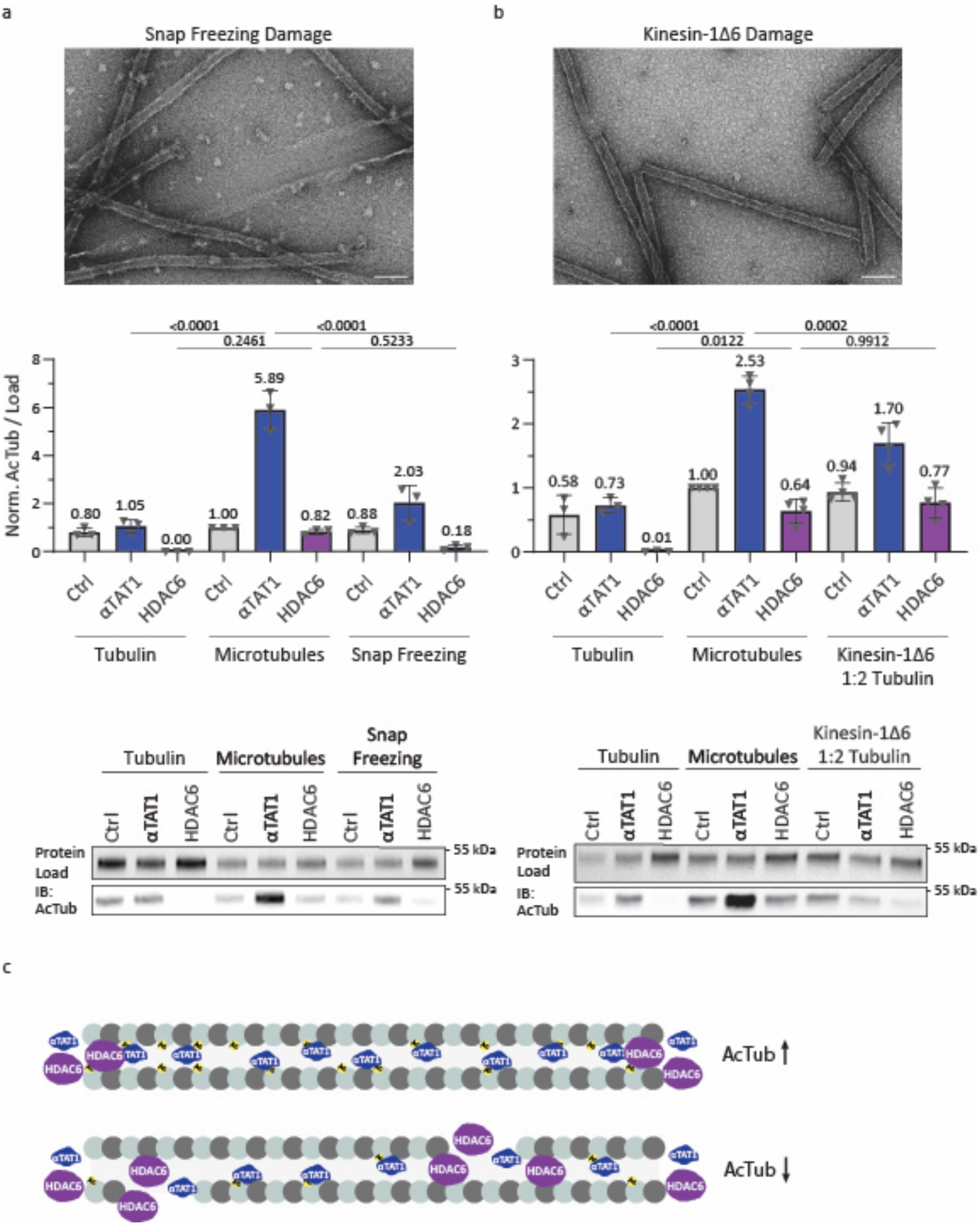
Microtubule damage reduces αTAT1 acetylation efficiency. **a** and **b** Representative EM images of microtubules (top) after (**a**) snap freezing or (**b**) kinesin-1Δ6-generated damage. Scale bars: 200 nm. Representative Western blot analysis and quantification (bottom) of normalized (Norm.) acetylation levels for both damage conditions compared to tubulin and intact microtubule as control conditions, for αTAT1, HDAC6 or in absence of the enzymes (Ctrl). Statistics: one-way ANOVA. Mean with SD. **c** Cartoon depicting acetylation levels in the presence of αTAT1 and HDAC6 of intact microtubules with increased acetylation (above) and damaged microtubules showing decreased acetylation (below). Without damage, αTAT1 diffuses through the microtubule and acetylates stochastically, while HDAC6 entry is limited. Damage along the shaft facilitates HDAC6 and αTAT1 access to the lumen, but αTAT1 acetylates only intact microtubules efficiently.

Due to its small size and rapid diffusion, αTAT1 equilibrates between the microtubule lumen and the cytosol within minutes in a 10-μm-long microtubule^49^. Detectable gradients or acetylation patches near damage would require microtubules longer than ∼150 μm^49^. Acetylation at K40 is therefore limited not by access to the lumen, but by the inherently slow catalytic rate of αTAT1. While αTAT1 can enter through damage sites, it does not rely on them to access K40^43^; entry through microtubule ends is sufficient.

Although microtubule damage can, in principle, grant luminal access, we found that microtubule damage does not enhance αTAT1’s effective activity - in fact, it impairs it. Kinesin-1Δ6-treated microtubules showed a 2-fold reduction in αTAT1-mediated acetylation compared to undamaged controls (Fig. 3b). This effect was further amplified by snap freezing, where damaged microtubules showed a 5-fold reduction in acetylation which accounts for only a 2-fold increase in acetylation, compared to a nearly 6-fold increase in intact microtubules (Fig. 3a). This unexpected decrease in αTAT1 efficiency correlates with the degree of lattice disruption. However, it does not scale with damage frequency - an increase of 0.3 events per micrometer is unlikely to explain a 2-fold reduction in acetylation. This suggests that damage sites exert long-range effects on the microtubule lattice that impair acetylation efficiency beyond the immediate vicinity of the defect. Given that αTAT1 binding depends on lateral tubulin interactions^49^, we propose that αTAT1 requires an intact and continuous microtubule lattice for effective recognition of the K40 loop and subsequent acetylation (Fig. 3c).

### αTAT1 efficiently acetylates expanded microtubule lattices

Microtubule structural states play a role in modulating interactions with microtubule-associated proteins (MAPs) - which can read out and/or induce lattice compaction or expansion - such as EB3, DCX^9,10^, TPX2^11^, MAP7 and tau^12,13^. We therefore asked whether αTAT1’s acetylation efficiency is influenced by specific microtubule lattice conformations.

To fine-tune lattice spacing, we polymerized microtubules under different conditions: in the presence of GMPCPP, GTP, a 1:1 mixture of GMPCPP and GTP, or taxol. In the presence of GMPCPP or taxol, microtubules are known to adopt an expanded lattice state (83.2 Å and 82.3 Å respectively)^4^. The 50:50 GMPCPP/GDP condition was designed to generate an intermediate lattice state, while GDP alone (after GTP hydrolysis) yields a compacted lattice (81.5 Å)^4^. When incubated with αTAT1 for 10 minutes, GMPCPP- and taxol-stabilized microtubules showed the highest acetylation levels, with a 4.59-fold and 4.20-fold increase, respectively (Fig. 4a). Notably, GMPCPP-microtubules - having the most expanded lattice - were slightly more acetylated than taxol-stabilized ones. The intermediate GMPCPP/GDP state showed less increase in acetylation with 3.05-fold, and a GDP-compacted microtubule lattice led to the lowest acetylation increase of only 1.80-fold.

**Fig. 4.**
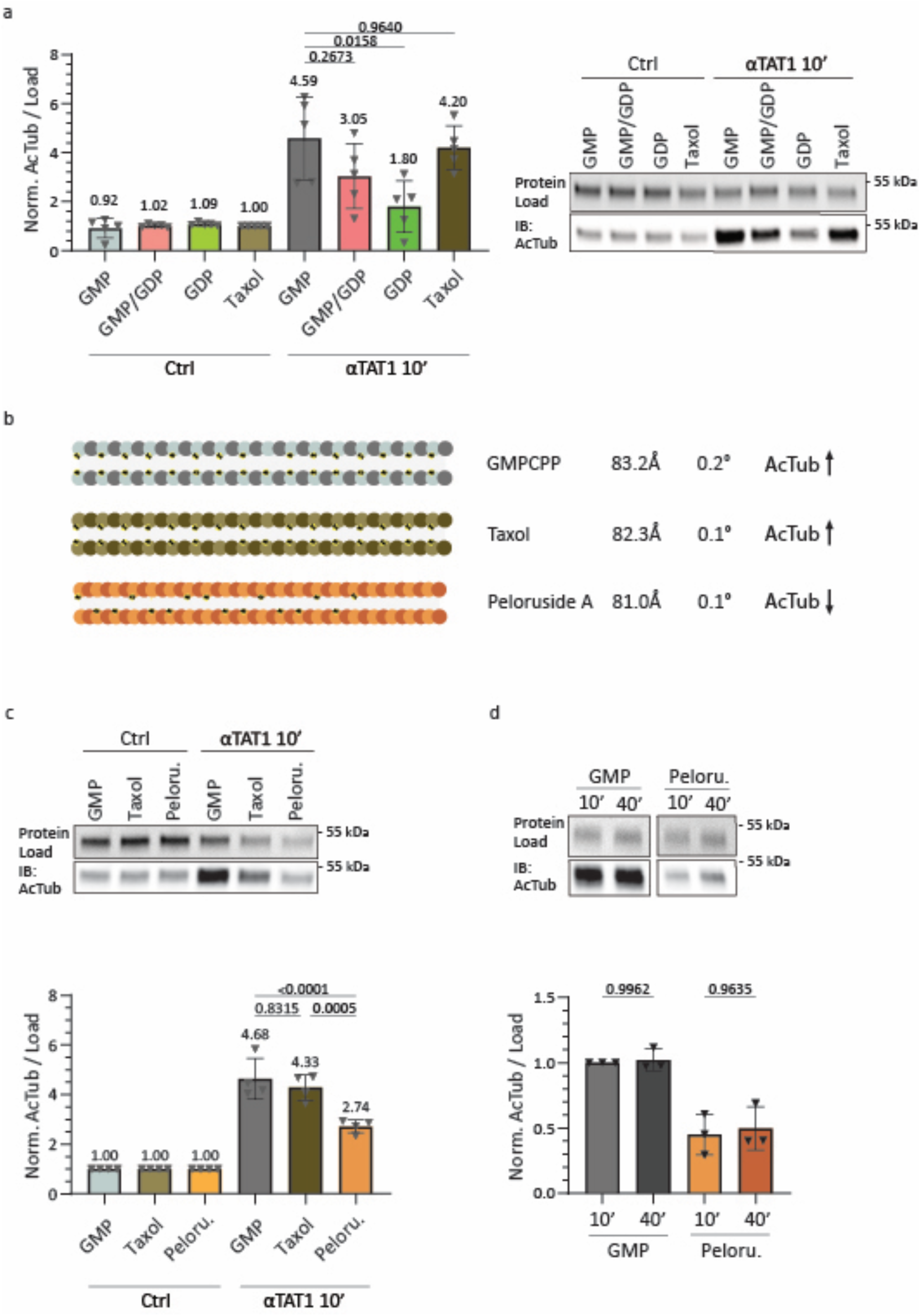
Lattice expansion and positive tubulin twist increase acetylation of microtubules. a, c and. **d** Representative Western blot analyses for microtubule acetylation reactions with αTAT1 for (**a**) 10 min in presence of GMPCPP, GMPCPP/GDP 1:1 ratio, GDP or taxol microtubules compared to control conditions; (**c**) 10 min (10’) on GMPCPP (GMP), Taxol or Peloruside A (Peloru.) stabilized microtubules compared to control conditions; Note, GDP-microtubules result from hydrolysis of GTP shortly after addition at the growing microtubule end. (**d**) 10 min or 40 min (40’) on GMPCPP- or Peloruside A- stabilized microtubules. Respective quantifications of AcTub levels relative to the protein load are plotted, from five (**a**), four (**c**) or three (**d**) independent experiments. Statistics: one-way ANOVA. Mean with SD. **b** Cartoons displaying different microtubule lattice conformations with GMPCPP, Taxol or Peloruside A. Dimer rise, and dimer twist values are indicated according to Kellogg et al.^5^ leading to a de-/ increase in microtubule acetylation levels indicated with arrows.

We expected that the microtubule lattice expands near damage sites, which - based on our findings with GMPCPP- and taxol-stabilized microtubules - should enhance acetylation by αTAT1. However, our results with kinesin-1Δ6–treated and snap-frozen microtubules contradict this assumption: instead of increased acetylation, we observed a reduction. One possible explanation is that tubulin does not expand near damage sites. Another one is that local, uneven protofilament expansion around damage sites disrupts lateral tubulin contacts, impairing αTAT1’s ability to recognize and modify the K40 loop. Unfortunately, we currently lack tools to resolve the tubulin and loop structure around a damage site at high resolution, so we relied on indirect, lattice-modulating assays as further readouts.

In addition to lattice compaction and expansion, different nucleotides and MAPs can alter the twist of tubulin dimers within the microtubule lattice. To distinguish the effects of lattice spacing from dimer twist, we tested another stabilizing drug: peloruside A. Unlike GMPCPP and taxol, peloruside A locks microtubules in a compacted lattice conformation (81.0 Å)^5^ when added during polymerization. It also increases the positive lattice twist by 10-fold, from 0.01° in GDP-microtubules to 0.1°^5^. Taxol, while expanding the lattice, induces a similar twist (∼0.1°), whereas GMPCPP-stabilized microtubules exhibit the highest positive twist (∼0.2°). We compared αTAT1 activity on microtubules stabilized with GMPCPP, taxol, or peloruside A (Fig. 4b). After 10 minutes of incubation, peloruside A–treated microtubules showed a 2.74-fold increase in acetylation (Fig. 4c) - higher than GDP-microtubules, which share a similar compacted lattice but have lower twists (Fig. 4b). However, peloruside A– stabilized microtubules were significantly less acetylated than taxol-stabilized ones, which showed a 4.33-fold increase despite having a comparable twist but a more expanded lattice (Fig. 4b,c). Extending the incubation time to 40 minutes did not substantially increase acetylation levels in peloruside A–treated microtubules, indicating that the observed differences in αTAT1 efficiency are not due to delayed kinetics but reflect persistent restrictions on catalytic activity (Fig. 4d). Taken together, these results show that while increased lattice twist may modestly enhance αTAT1 activity, lattice expansion has a greater impact on its enzymatic efficiency.

## DISCUSSION

Microtubule shaft damage can be induced by mechanical or physiological cues. In our reconstituted system we artificially introduce mechanical damage through repeated freezing and thawing cycles of microtubules (snap freezing). We address physiological damage by using a highly damaging mutant of kinesin-1 (kinesin-1Δ6) to reinforce motor protein-induced damage. In both cases, the stress exerted on microtubules leads to tubulin dimer dissociation from the lattice leaving behind a void within the polymer (Fig. 2). Motor stepping along the microtubule shaft most likely prompt local dissociation of multiple dimers within and across protofilaments which results in appearance of “kink” formations along the microtubule. Enhanced motor activity for high kinesin-1Δ6 concentrations promotes locally increased dimer dissociation resulting in breakage of the tube, referred to as “broken MTs” (Fig. 2a, d-f).

The enzymes αTAT1 and HDAC6 require access to the microtubule lumen for modification of the α- tubulin K40 residue. We and others showed that HDAC6 deacetylates non-polymerized tubulin dimers more efficiently than polymerized microtubules, most likely due to its size that limits its entry and diffusion in the microtubule lumen^54^ (Fig. 1b). The catalytic domain of HDAC6 comprises an area of 4x10 nm according to alphafold predictions (AF-Q9UBN7) which is marginally smaller than the 14 nm inner diameter of a microtubule^61^. Upon pronounced microtubule damage, we hypothesize that HDAC6 can reach the K40 loop more easily, and deacetylation therefore increases (Fig. 3a). While small damages encompassing only few dimers seem to be not sufficient to grant access (Fig. 3b). These results are in line with previous findings on higher HDAC6 deacetylation efficiency on polymerized Zn sheets or ”inside-out” tubulin-dolastatin rings compared to intact microtubules^54^.

αTAT1-mediated acetylation on the other hand increases more than 5-fold for polymerized microtubules (Fig. 1b,c). The structured region of αTAT1 comprises an area of only 3x5 nm (PDB: 4GS4). Hence, the rather small enzyme can enter through microtubule ends and freely diffuse through the lumen to modify the K40 residue of a 10 μm microtubule within 8 min^49^. αTAT1 efficiency is therefore dictated by its slow enzymatic activity rather than its access to the microtubule lumen^49^. Nevertheless, shaft damage induced by snap freezing of microtubules or by incubating microtubules with kinesin-1Δ6 motor proteins drastically decreases enzymatic efficiency for αTAT1 by up to 65% (Fig. 3).

We propose that αTAT1 is sensitive to microtubule lattice integrity. Any kind of damage to the microtubule shaft may alter the conformation of tubulin dimers in the lattice near the damage site, interfering with αTAT1’s ability to recognize the K40 loop. Our results demonstrate that αTAT1 can detect microtubule conformation ranging from compacted (81.0 Å) to expanded (83.2 Å) lattice states (Fig. 4) and responds accordingly. Furthermore, αTAT1’s ability for acetylation is sensitive towards the twist of the lattice. Hereby, the detected changes in acetylation efficiency are independent from nucleotide or protofilament number. For example, GMPCPP- and taxol-stabilized microtubules - despite having 14 and 13 protofilaments, respectively - are acetylated to similar levels, indicating that protofilament number does not influence αTAT1 activity under these conditions. However, acetylation efficiency decreases by half when microtubules are in a compacted state compared to expanded lattices (Fig. 4a,c,d). To acetylate, αTAT1 needs to recognize the K40 loop with its active binding site to transfer the acetyl group from Acetyl-CoenzymeA (AcCoA) to the lysine40 of α-tubulin, whereby the lateral contact to neighboring microtubules is important^49,62^. We suggest that a compaction of tubulin dimers limits access to the K40 loop exposed to the microtubule lumen and therefore reduces the efficiency of αTAT1 acetylation.

Our findings show that αTAT1 can detect microtubule lattice conformation *in vitro*, with a strong preference for acetylating microtubules that are intact and in an expanded lattice state. Since MAPs can modulate microtubule lattice spacing^8,11,12,16^, such regulation could influence where and how efficiently αTAT1 acts within cells. Indeed, it was shown that MAP7 which binds to the outside of the microtubule enhances αTAT1 acetylation efficiency^63^. Acetylation could specifically mark stable microtubules that adopt an expanded lattice state. How these conformational changes contribute to the spatial organization of acetylation patterns - such as the acetylation gradients observed with its distinct acetylation segments along microtubules in cells - remains an open question. Previously, we reported that acetylation distributes in a gradient in cells with high perinuclear acetylation that decreases towards the cell periphery and that running motor proteins modulate the gradient through microtubule damage^43^. Local damage caused by motor proteins could be a general mechanism to control lattice conformation of a subset of microtubules together with MAPs regulating the lattice spacing. Tubulin modifying enzymes like αTAT1 are sensitive to such structural changes of the microtubule shaft, leading to spatially patterned posttranslational modifications of the microtubule network in cells.

## METHODS

### Plasmid generation

From pEF5B-FRT-GFP-αTAT1(D157N) (addgene #27100), the amino acid at position 157 was mutated to aspartate and the GFP sequence was replaced by a Flag-tag using AgeI/BglII restriction enzymes. Flag-αTAT1 was subcloned into a pET29b vector for bacterial protein expression using KpnI/BamHI restriction enzymes. The highly damaging constitutively active kinesin-1 mutant (kinesin-1Δ6-GFP) was generated as previously described^43^ and cloned into pET17-K560-GFP-His (addgene #15219) using EcoRI/AgeI restriction enzymes. The HDAC6 sequence from HDAC6-Flag (addgene #13823) was subcloned into a pFastBachHTA vector using PluTI/NotI restriction enzymes for Baculovirus generation and subsequent Sf9 cell infection.

### Tubulin purification and labeling

Tubulin was purified from bovine brain by two polymerization/depolymerization cycles in High-Molarity PIPES buffer as described previously^64^. In brief, a first polymerization/depolymerization was performed in High-Molarity PIPES buffer (1 M PIPES pH 6.9, 10 mM MgCl2, 20 mM EGTA, 1.5 mM ATP, 0.5 mM GTP) supplemented with 1:1 glycerol and Depolymerization buffer (50 mM MES at pH 6.6, 1 mM CaCl2) respectively. A second polymerization/depolymerization was performed with polymerization in High-Molarity PIPES buffer and depolymerization in 0.25x BRRB80 with addition of 5x BRB80 after 15 min to reach 1xBRB80 (80 mM PIPES at pH 6.8, 1 mM MgCl2, 1 mM EGTA).

Purified tubulin was labeled with ATTO-488, ATTO-565, ATTO-647 (ATTO-TEC GmbH) or biotinylated tubulin, adapted from a previously published protocol^65^. Briefly, tubulin was polymerized in glycerol PB solution (80 mM PIPES pH 6.8, 5 mM MgCl2, 1 mM EGTA, 1 mM GTP, 33% glycerol) for 30 min at 37 °C and placed in layers onto cushions of 0.1 M NaHEPES pH 8.6, 1 mM MgCl2, 1 mM EGTA and 60% glycerol and centrifuged. The pellet was resuspended in resuspension buffer (0.1 M NaHEPES pH 8.6, 1 mM MgCl2, 1 mM EGTA and 40% glycerol) followed by incubation for 10 min at 37 °C with either 1/10 volume of 100 mM ATTO-488, -565, or -647-NHS-fluorochromes or for 20 min at 37 °C with 2 mM NHS-biotin. Labeled tubulin was sedimented onto cushions of 1x BRB80 supplemented with 60% glycerol, resuspended in BRB, and a second polymerization/depolymerization cycle was performed. The labeling ratio was 11% for ATTO-488 and 12% for ATTO-565.

### Protein expression and purification

For purification of <u>αTAT1</u>, *E.coli* BL21 (DE3 pLysS) cells were transformed and grown at 37 °C until an optical density at 600 nm of 0.6-0.8 and induced for protein expression with 1 mM IPTG at 18 °C overnight. Cells were harvested and lysed in lysis buffer (50 mM Tris pH 7.5, 150 mM NaCl, 10% glycerol) supplemented with 1% Triton-X 100, 2 mM DTT, 2 mM PMSF and protein inhibitor cocktail (Roche) and subsequently sonicated. Lysed cells were cleared by centrifugation at 12000 rpm for 30 min at 4 °C. Proteins were affinity purified using a 1mL HisTrap HP column (GE Healthcare), washed in lysis buffer supplemented with 0.2% Triton-X 100 and 25 mM imidazole and eluted with 250 mM imidazole within 10 column volumes. Fractions were pooled and concentrated using Amicon 30K Centrifugal Filters (Millipore). Concentrated proteins were loaded onto a size-exclusion chromatography using a HiLoad 16/600 Superdex 200 column (GE Healthcare) in 20 mM Tris pH 7.5, 150 mM NaCl, 1 mM MgCl2. Collected fractions were concentrated and concentration determined by Bradford assay. Proteins were supplemented with 10% glycerol, aliquoted, snap-frozen in liquid nitrogen and stored at -80 °C.

For <u>kinesin-1Δ6-GFP</u> (referred to as: kinesin-1Δ6) a similar protein purification protocol was used. Protein expression was induced with 0.5 mM IPTG at 16 °C overnight. Cells were lysed in 100 mM Na2HPO4/NaH2PO4 pH 7.5, 600 mM KCl, 3 mM MgCl2, 10% glycerol, 20 mM β-mercaptoethanol, supplemented with 10 mM imidazole, protein inhibitor cocktail and 100 mM ATP addition right before lysis. Cell lysates were ultracentrifuged at 40000 rpm for 50 min at 4 °C. Proteins were loaded on a 1 mL HisTrap HP column were washed with 30 mM imidazole, washed again with 60 mM and eluted with a 60-500 mM imidazole gradient over 30 column volumes. Size-exclusion chromatography was performed in 10 mM HEPES pH 7.4, 150 mM KCl, 1 mM MgCl2, 0.05 mM ATP, 1 mM DTT and concentrated proteins were supplemented with 20% sucrose before snap-freezing and storage at - 80 °C.

<u>HDAC6</u> was purified from insect cells by resuspending the cell pellet in lysis buffer (100 mM Tris pH 8, 10 mM NaCl, 5 mM KCl, 2 mM MgCl2, 10% glycerol, 0.2% Triton-X) supplemented with protein cocktail inhibitor. After lysis, NaCl concentration was adjusted to 150 mM, and cells were ultracentrifuged at 50000 rpm for 20 min at 4 °C. Affinity chromatography was performed using a 1 mL HisTrap HP column and proteins were washed with 25 mM imidazole and eluted from the column with 250 mM imidazole within 5 column volumes. Concentrated proteins were further purified by size-exclusion chromatography in lysis buffer and concentrated proteins were supplemented with 10% glycerol, snap-frozen and stored at -80 °C.

### Microtubule seed preparation

Short microtubules seeds were prepared by mixing 30% ATTO-647-labeled tubulin with 70% biotinylated tubulin with a final concentration of 10 μM in 1x BRB80. The solution was supplemented with 0.5 mM GMPCPP and incubated for 45 min at 37 °C. After the addition of 1 μM Paclitaxel (Taxol), the solution was incubated again for 30 min at 37 °C and subsequently centrifuged for 15 min at 14000 rpm at room temperature (RT). The microtubule pellet was resuspended in 1x BRB80 supplemented with 0.5 mM GMPCPP and 1 μM Taxol. Seeds were aliquoted and stored in liquid nitrogen.

### Microtubule assembly

GMPCPP-stabilized microtubules were assembled by incubating 5 μM ATTO-488 or ATTO-565 labeled tubulin in 1x BRB80 buffer (80 mM PIPES pH 6.8, 1 mM MgCl2, 1 mM EGTA) supplemented with 0.5 mM GMPCPP for 20 min at 37 °C, followed by two subsequent additions of 1 μM tubulin for 10 min at 37 °C. The tubulin mixture was centrifuged at 12000 rpm for 15 min at RT and microtubules resuspended in 1x BRB80. For comparison with other stabilized microtubule lattices, GMPCPP microtubules were assembled from 10 μM directly without sequential tubulin addition.

### Microtubule (de)acetylation

GMPCPP-stabilized microtubules (1 μM tubulin) were diluted in 1xBRB supplemented with 100 μM Acetyl CoenzymeA (Sigma-Aldrich, A2056) and 1 mM DTT. For (de)acetylation reactions with equimolar ratios, 1 μM αTAT1, 1 μM HDAC6 or the same volume of HDAC6 purification buffer as a control were added to different tubes respectively. For enzyme competition experiments, 1 μM αTAT1 and 1 μM HDAC6 were added to the same tube or either αTAT1 or HDAC6 concentrations were lowered by 10-fold to 0.1 μM. As a control for microtubule acetylation, the same procedure was performed with non-polymerized tubulin (1 μM). Reactions were incubated for 1h at RT and samples taken for Western blotting or immunostaining (for microtubules only).

### Microtubule acetylation assay for different lattices

For generating microtubules with different microtubule lattices, 10 μM ATTO-488 or ATTO-565 labeled tubulin was incubated with 0.2 μM microtubule seeds in 1x BRB80 buffer supplemented with 1 mM GMPCPP, 0.5 mM GMPCPP/0.5 mM GTP, 1 mM GTP or 1 mM GTP/10 μM taxol respectively.

Taxol-, or peloruside A-stabilized microtubules were assembled similarly to GMPCPP-stabilized microtubules from 10 μM ATTO-488 or ATTO-565 labeled tubulin in 1x BRB80 supplemented with 1 mM GTP and 1 μM taxol or 1 μM peloruside A (a kind gift from Michel Steinmetz, Peter Northcote and John Miller) respectively. Microtubules were polymerized for 30 min at 37 °C and 10 μM αTAT1 with 100 μM Acetyl Coenzyme A was subsequently added to each mixture and incubated for 10 min at 37 °C. Reactions were centrifuged at 12000 rpm for 15 min at RT and microtubules resuspended in 1x BRB. Samples were taken for Western blotting or immunostaining.

### Microtubule damage generation

GMPCPP-stabilized microtubules (7 μM tubulin) were exposed to five subsequent cycles of snap freezing in liquid nitrogen for 1 min, followed by thawing the solution back to RT for 1 min. After completion of freezing/thawing cycles, microtubules were immediately stained for electron microscopy or used for (de)acetylation assays. For control conditions, untreated GMPCPP-microtubules were used.

For damaging microtubules with kinesin-1Δ6, GMPCPP-stabilized microtubules (3.5 μM tubulin) were incubated with either 1.75 μM or 84 nM of kinesin-1Δ6 for a 1:2 or 1:24 tubulin ratio respectively, in 1x BRB supplemented with 2.7 mM ATP. Control conditions were performed in 1x BRB with 2.7 mM ATP only. Reactions were incubated for 15 min at 37 °C and immediately used for further assessment.

### Microtubule (de)acetylation of damaged microtubules

Intact, snap freezing- or kinesin-1Δ6-damaged microtubules (1 μM tubulin) or non-polymerized tubulin (1 μM) were diluted in 1xBRB supplemented with 100 μM Acetyl CoenzymeA and 1 mM DTT. For (de)acetylation reactions, 1 μM αTAT1 or 1 μM HDAC6 or the same volume of HDAC6 purification buffer as a control were added respectively. Reactions were incubated for 1 h at RT and samples taken for Western blotting.

### Negative stain transmission electron microscopy

For negative stain transmission electron microscopy (negative stain EM), damaged microtubules were diluted in 1x BRB to a final concentration of 3.5 µM. A 4 µL drop of the sample was applied directly onto glow-discharged copper EM grids (Electron Microscopy Sciences). After a one-minute incubation, the excess sample was blotted off, and the grids were washed twice with 10 µL drops of 2% uranyl acetate (Electron Microscopy Sciences), each time followed by immediate blotting. The grids were then stained with a 10 µL drop of 2% uranyl acetate for one minute, blotted, and air-dried for two minutes. Prepared grids were stored in EM grid boxes at RT until imaging. Imaging was performed using a Talos L120C transmission electron microscope.

### SDS-PAGE and Western blot

Samples were diluted in 1x sample buffer (60 mM Tris, 2% SDS, 10% glycerol, 5% β-mercaptoethanol, 0.005% bromphenol blue) and run on 4-20% Mini-PROTEAN® TGX Stain-Free™ Protein Gels (BIO-RAD) in 1x running buffer (25 mM Tris, 192 mM Glycine, 0.1% SDS, pH 8.3). Protein load was detected by UV exposure of gels with a Fusion FX Vilber (Witec AG). Proteins were transferred to a nitrocellulose (NC) membrane of iBlot™ 3 Transfer Stacks (ThermoFisher Scientific) with an iBlot™ 3 Western Blot Transfer Device (ThermoFisher Scientific). NC membranes were blocked for 1 h with blocking buffer (5% dried milk in TBS-Tween 0.5%) and incubated with the primary antibody anti-acetylated tubulin (Sigma-Aldrich, T74451, 1:1000 dilution) in blocking buffer at 4 °C overnight. The next day, membranes were washed 3x with TBS-Tween 0.5% (TBS-T) and incubated for 1 h at RT with the secondary antibody anti-mouse-horseradish peroxidase (HRP) (GE Healthcare, NA9310, 1:5000 dilution) in blocking buffer. Membranes were washed again 3x with TBS-T and HRP-conjugated proteins were revealed with a WesternBright® ECL detection kit (Advansta) on a Fusion Solo Vilber Lourmat camera (Witec AG). The UV protein load signal was used as a loading control for normalization of the anti-acetylated tubulin signal. The values for the acetylated tubulin antibody signal / UV protein load were normalized to a control condition in each experiment.

### Immunofluorescence

Coverslips were cleaned in two steps: sonication in 1 M NaOH for 40 min, rinsing in bidistilled water, followed by sonication in 96% ethanol for 40 min, repeated rinsing in bidistilled water and then dried with an air gun. *In vitro* polymerized microtubules were sedimented for 5 min on cleaned coverslips and subsequently fixed with 100% cold methanol for 4 min at -20 °C, followed by 3x washes with phosphate buffered saline (PBS). Microtubules were permeabilized and blocked with blocking buffer [2% bovine serum albumin (BSA) and 0.1% Triton X-100 in PBS] for 5 min and subsequently incubated with the primary antibodies anti-acetylated tubulin (Sigma-Aldrich, T74451, mouse, 1:1000 dilution) or anti-α-tubulin (Geneva Antibody Facility, AA345, rabbit, 1:1000 dilution) in blocking buffer in a closed humidified chamber at RT for 15min. Microtubules were washed 3x with blocking buffer and incubated with the secondary antibodies (Invitrogen, species-specific IgG conjugated to Alexa-647, or 488 fluorophores, 1:1000 dilution in blocking solution) for another 15 min in a closed humidified chamber at RT. Microtubules were washed again 3x with blocking buffer and coverslips were mounted onto glass microscopy slides (Glass technology) and sealed with nail polish. Immunofluorescence images were taken using an Axio Observer Inverted TIRF microscope (Zeiss, 3i) and a Prime 95B (Photometrics). A 100X objective (Zeiss, Plan-Apochromat 100X/1.46 oil DIC (UV) VIS-IR). SlideBook 6X64 software (version 6.0.4) was used for image acquisition.

### Image and statistical analysis

Immunofluorescence images were analyzed using ImageJ. For data analysis a background of 100 a.u. was subtracted for each channel.

The <u>average acetylation level</u> of reconstituted microtubules was determined by measuring the fluorescence intensity profile of the acetylated tubulin antibody relative to the fluorescence intensity profile of the microtubule polymerized from purified and ATTO-488- or ATTO-565-labeled bovine brain tubulin (as described here above) and normalized to the control condition. For the <u>acetylation profile</u> of different subsets of microtubules, the intensity values for acetylated tubulin / ATTO-tubulin were Ln (Natural Log) transformed and plotted over the normalized distance. To normalize each microtubule, the length of the line plot was divided by the maximum value. For comparison of individual microtubules, the normalized distance values were binned into 0.005 steps and the combined acetylation profiles averaged.

For electron microscopy analysis in ImageJ, the total microtubule length was measured in each image and summed for all conditions individually. The event count was divided by the total microtubule length determined for each condition. The percentage of opened microtubule sheets was calculated by summing and dividing the length of all detected sheet openings per condition by the total microtubule length.

Statistical analyses were performed by one-way ANOVA with Tukey’s multiple comparisons test. using GraphPad Prism software v.10.4.1. P values less than 0.05 were considered statistically significant.

## DATA AVAILABILITY

All data associated with this study are presented in the manuscript in main figures and the supplementary information.

## ACKNOWLEDGEMENTS

CE has been supported by the SNSF, (TMSGI3_211433), JT has been supported by the DIP of the Canton of Geneva. CA has been supported by the DIP of the Canton of Geneva, SNSF (310030_212563, CRSII5_216597 and TMSGI3_211433).

## AUTHOR CONTRIBUTIONS

CE and CA conceptualized the study and designed the experiments. CE performed and analyzed all experiments. MCV generated the plasmids used in this study. JT performed electron microscopy and CE analyzed the images. CE and CA wrote the manuscript.

## LIMITATIONS OF THE STUDY

A limitation in the study is the widely used, commercially available antibody for tubulin acetylation 6- 11B-1 (Sigma-Aldrich, T74451) for *in vitro* assays as previously reported. As described, the epitope recognition is perturbed in the presence of the microtubule stabilizing agent taxol^46^ and likely by other factors impacting the lattice structure (Fig. S1c). Consequently, imaging analysis of immunofluorescence labeled microtubules *in vitro* is restricted.

---

**Fig. S1.**
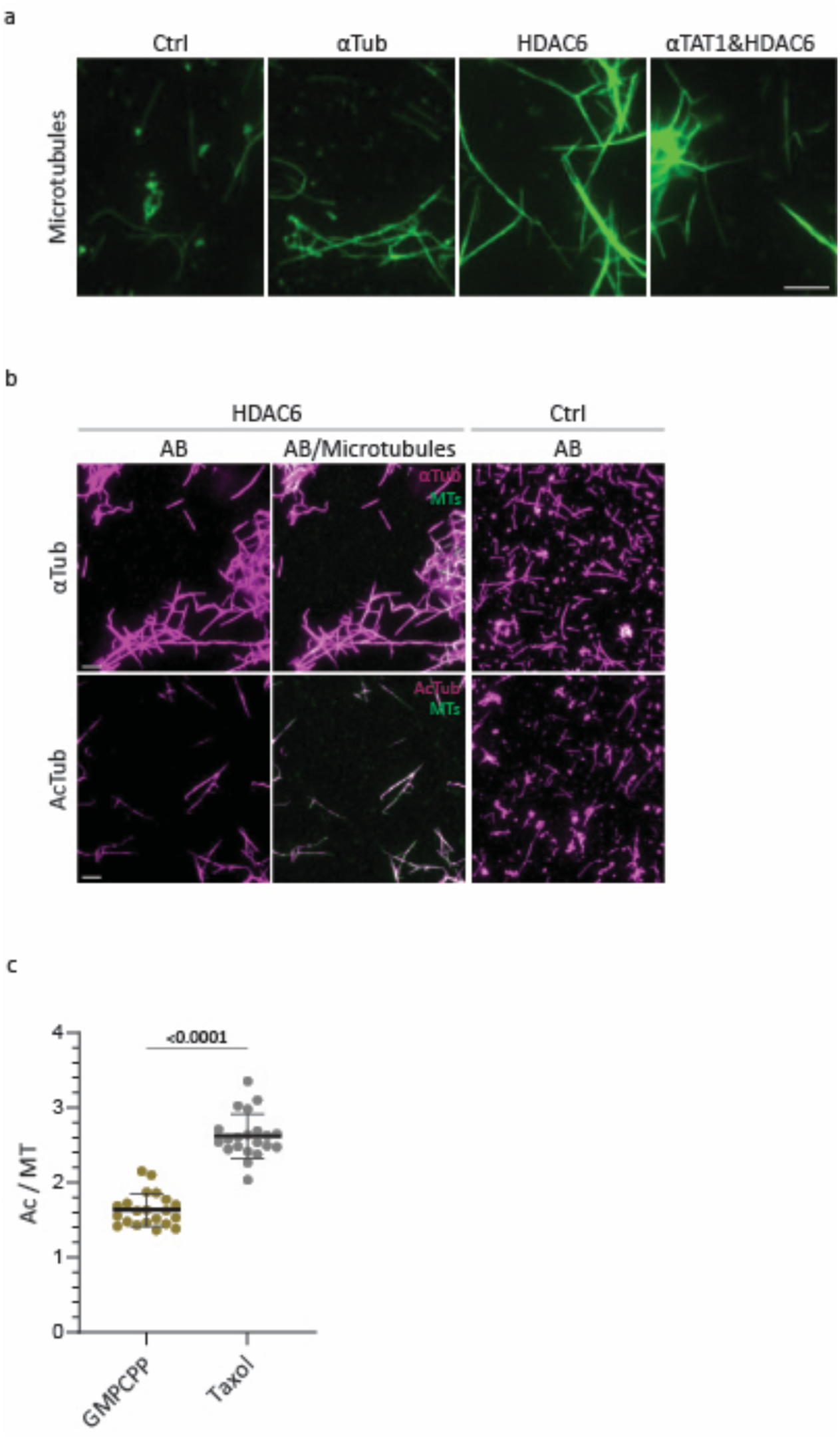
Taxol affects Ac-K40 epitope recognition by Antibody. **a** and **b** Representative immunofluorescence images of fluorescent microtubules (green) that were incubated with (**a**) αTAT1, HDAC6, both enzymes or control conditions and fixed; or (**b**) with HDAC6 or control conditions, fixed and immunostained with alpha-tubulin (αTub) or acetylated tubulin (AcTub) antibodies (AB) (magenta). Scale bars: 5 µm. **c** Immunofluorescence signals of acetylated tubulin over microtubules for GMPCPP (n=21) or taxol (n=21) in control conditions from one experiment. Statistics: two tailed t test. Mean with SD.

**Fig. S2.**
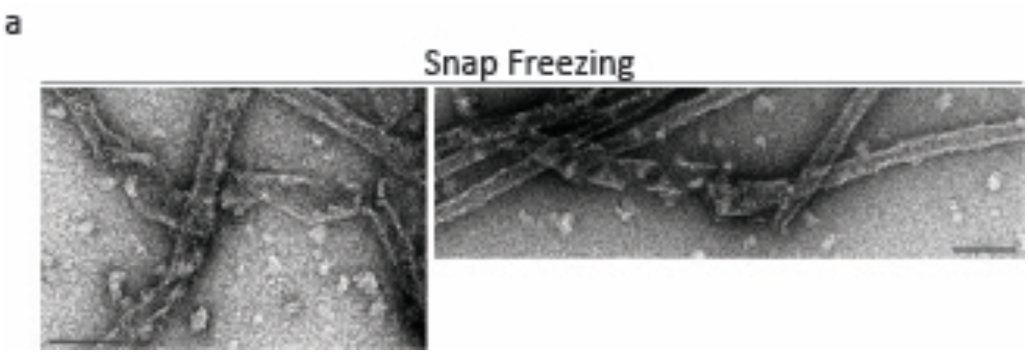
Distinct damage phenotype for Snap Freezing microtubule damaging conditions. **a** Representative EM images for snap freezing conditions with “curled” sheet opening phenotype. Scale bars: 100 nm.

